# Evolutionary impact of *in vitro* adaptation on virulence in the pathogen *Zymoseptoria tritici*

**DOI:** 10.1101/2023.12.31.573786

**Authors:** A. J. Jallet, G. Robert-Siegwald, A. Genissel

**Affiliations:** Université Paris Saclay, INRAE, UR BIOGER, 91120, Palaiseau, France

**Keywords:** abiotic environment, genome integrity, *de novo* mutation, plant-pathogen evolution, secretome

## Abstract

- All species are living in variable environments. integrating the impact of changing environments into plant-pathogen studies becomes essential. This integration is key to expanding our understanding of the evolutionary dynamics governing plant-pathogen interactions.
- After subjecting Zymoseptoria tritici to 500 generations of experimental evolution in vitro under various temperature regimes, we assessed the evolved lineages’ virulence levels across six wheat cultivars. Additionally, we performed whole-genome sequencing on ten evolved lineages to identify accumulated mutations
- Our study revealed a reduction in virulence among several evolved lineages, with variability based on the host genotype. We observed trans-lineage segregating mutations in the genome, encompassing both synonymous and non-synonymous mutations within the secretome. Intriguingly, despite 500 generations of experimental evolution, no loss of dispensable chromosomes was detected
- These findings suggest that the abiotic environment can significantly influence the dynamic evolution of the plant pathogen *Z. tritici*.

## Introduction

Understanding how species are able to respond to environmental variations remains a challenge in the field of Evolutionary Biology. In the quest to understanding the dynamic arms race between plants and their pathogens, the impact of changing environments has been rarely considered, despite its widespread influence on plant-pathogen evolution (Velasquez et al., 2018). Studies in laboratory controlled conditions hence represent a first step towards unravelling the complexity of plant pathogens within the framework of global change. In the present study our model is the ascomycete fungus *Zymoseptoria tritici,* a pathogen responsible for Septoria Tritici Blotch (STB), one of the most important foliar diseases of wheat (Fones and Gurr, 2015; O’Driscoll et al., 2014).

Our understanding of the intricate molecular interactions between wheat and this fungal pathogen is continually expanding (Kettles and Kanyuka, 2016). Currently, there are approximately 21 recognized resistance genes, along with numerous other Quantitative Trait Loci (QTLs) that breeders consistently incorporate into wheat cultivars to enhance resistance (Brown et al., 2015; Saintenac et al., 2021), but investigations on the genetic basis of fungal pathogen virulence have been largely neglected (Bartoli and Roux, 2017; Genissel 2017). There has been no study on the genetic basis underlying virulence of *Z. tritici* but a few recent ones, advanced by NGS technologies which open new opportunities for non-model species. Since the publication of the first complete genome of *Z. tritici* (Goodwin et al., 2011), two main approaches are thus employed to dissect the genotype-phenotype link for the quantitative variation of virulence among *Z. tritici* populations: bi-parental QTL mapping using RADseq and Association Genetics using whole genome sequencing. Two QTL mapping studies have been published thus far, revealing QTLs (or genes) of large effect on chromosome 7 (Stewart et al., 2017; Meile et al., 2018). Genome-Wide Association Studies (GWAS) using a sample size > 100 individuals have also identified two effector proteins (gene Avrstb6 on chromosome 5 by Zhong et al., 2017; Hartmann et al., 2017). It is now becoming quite clear from these studies that small secreted proteins are key components in the hide-and-seek evolutionary game between the pathogen and its host. In addition, each of these candidate effectors resides in proximity to transposable elements (TE), within genomic regions abundant in AT nucleotides. This might suggests that chromosomal rearrangements caused by TE are setting up the stage for rapid mutation accumulation in functionally relevant genes. Indeed despite the lack of evidence for their contribution to phenotypic variation many other effector genes are fast evolving and are located in regions with a high recombination rate (Poppe et al., 2015; Croll et al., 2015).

In this study, we present the virulence levels of lab-evolved populations originating from a single clone subsequent to 1 year *in vitro* evolution. We aimed to examine the hypothesis that one year of in vitro evolution could impact the pathogen’s capacity to invade its host. A previous study has indicated extensive transcriptome rewiring within the evolved strains, identifying candidate genes whose transcription levels might be implicated in adapting to temperature fluctuations. Furthermore, our observations revealed greater gene expression evolution at the tips of core chromosomes, as well as on the accessory chromosomes. In this study, we employed Next-Generation Sequencing (NGS) to investigate the mutations accumulated throughout the evolutionary process, aiming to establish their potential association with the observed phenotypic changes. The disease levels of 18 evolved strains and the two founder clones (referred to as MGGP01 and MGGP44) were evaluated across six bread wheat cultivars, each harboring different resistance genes. The key findings emphasize: (i) a higher occurrence of symptom loss in lineages evolving under fluctuating selection, (ii) the retention of accessory chromosomes throughout the experimental evolution, and (iii) the identification of multiple mutations in evolved lineages, leading to amino acid alterations, notably within effector genes.

## Materials and methods

### Biological material

Z. tritici strains used in this study resulted from an experimental evolution, testing for three different temperature regimes: stable at 17°C, stable at 23°C and a fluctuating regime between these two temperatures. Evolved lineages and ancestral clones used in this study are described in Figure 1. The two ancestors correspond to the strains MGGP01 and MGGP44 isolated in 2010 from lesions on cultivar Caphorn and cultivar Apache, respectively. Both strains were collected at the same location (South of France, Auzeville-Tolosane) (43° 53’ N - 1° 48’ W). Each strain was stored in 30% glycerol at −80°C. For sequencing, 10 strains were used (both ancestors and four evolved lineages derived from each, see **Figure 1**). Since the experimental evolution was conducted at two different temperatures (17°C and 23°C), cell multiplication for NGS was performed at 17°C and 23°C, in 500mL of liquid Potato Dextrose Broth medium, for each of the 10 strains, in the same conditions as for the experimental evolution (Jallet et al., 2019). Prior to phenotyping, aliquots from −80°C stocks were multiplied on petri dishes using Potato Dextrose Agar medium, at 20°C. For *in planta* experiments we used six bread wheat cultivars which are known to manifest different resistance level against STB in the French fields: Caphorn, Apache, Premio, Courtot, Obelisk and Taichung29 (**Figure 2A**).

**Figure 1 -.**
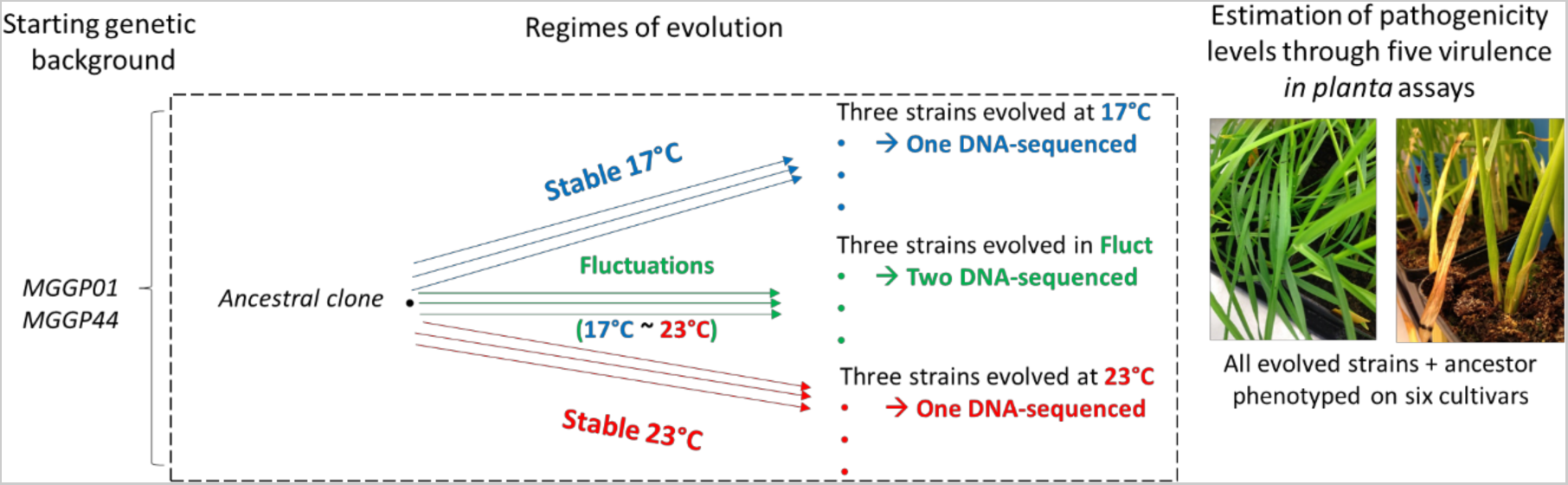
Experimental evolution design prior to in planta phenotyping. From each of the two clonal ancestors (MGGP01 and MGGP44), three replicates per regime evolved in the following conditions: at 17°C, at 23°C and under temperature fluctuations. All evolved lines plus the two ancestors were phenotyped on six wheat cultivars by measuring their ability to cause symptoms *in planta*. In addition, the two ancestors and four evolved lineages from each of them (one Stable_17°C, one Stable_23°C and two fluctuating) were whole-genome sequenced.

**Figure 2 -.**
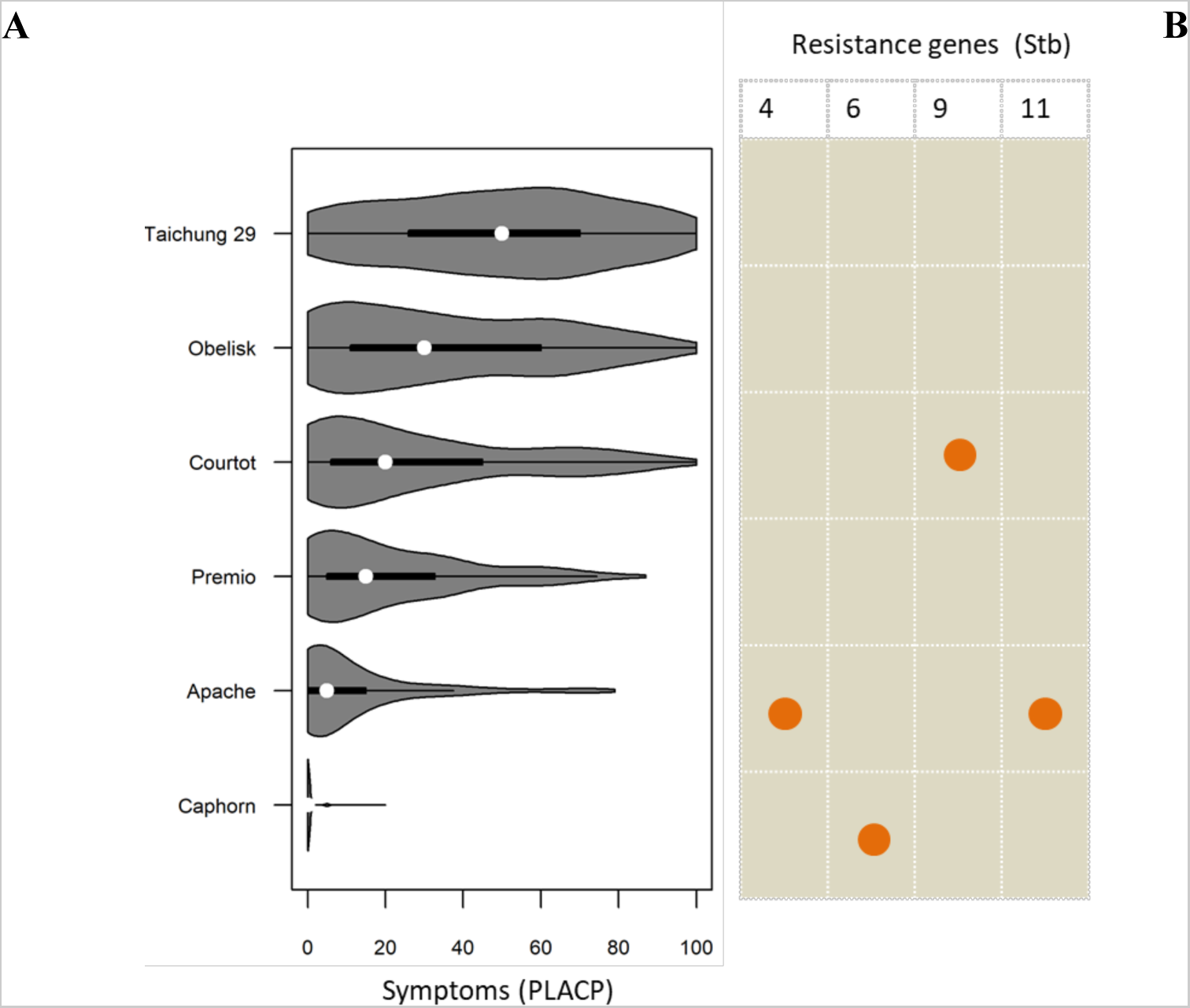
Overall disease symptom distribution for the six bread wheat cultivars. **A.** Overall distribution across all fungal genotypes (2 ancestors and 18 evolved lineages). **B.** Orange dots indicate the presence of known resistance genes (Stb genes) in the genome of each wheat cultivar (Taichung29, Obelisk, Courtot, Premio, Apache, Caphorn).

### Plant infection

Six days prior to inoculation fungal samples were multiplied in vitro on PDA medium at 20°C. Spores were diluted to a concentration of 10^7^ cells per mL in distilled water, using repeated counting with microglass chambers (Kova Glasstic R Slide 10). Tween 20 was added to the cell suspension to favour the fungal adherence at the leaf surface (0.1% v/v). For quality leaf development assessment and to ensure the absence of cross-contamination of fungal cell suspensions between leaves, a negative control (a leaf inoculated with water) was included in each pot. We planted 5 seeds, negative control included, per fungal genotype for each experiment. Five independent replicated experiments were conducted, thus totalizing 25 plants for each plant genotype-by-fungal genotype interaction. Before and after inoculation, plants were maintained in a large growth chamber (16h/8h day/night, 22°C/18°C, 80%/95% relative humidity). Leaf inoculation was performed on 14 days-old seedlings by painting cell suspensions onto a segment of 7 cm of the first true leaf. To enable fungal dispersion at the leaf surface, plants were maintained for 2.5 days at a higher humidity rate of 100%. At 12 days post inoculation (dpi), plants were watered with nutritive solution (*PlantProd* fertilizer). At 21 dpi, we monitored the quantitative variation of symptoms by visual inspection using binoculars. For each inoculated leaf we annotated the percentage of leaf area covered with lesions (pycnidia) (PLACL).

### Genomic DNA extraction

DNA was isolated for all *Z. tritici* lineages (**Figure 1**). Since the experimental evolution was conducted at two different temperatures, cell suspension and whole genome sequencing were also conducted at 17°C and 23°C. This design enables the examination of temperature-associated clone interference within the evolved lineage. Snap frozen lyophilized mycelium was grinded and vortexed in 14mL Lysis buffer at 65°C and then incubated for 30 minutes at 65°C. Samples were gently mixed in 3M Sodium Acetate pH5.2, incubated on ice for 30min and centrifuged at 10,000G for 15min at 4°C. Supernatant was transferred onto 0.7 volume of cold absolute isopropanol. Samples were centrifuged at 6,000G for 10min at 4°C, and pellets were washed in 4mL of cold 70% ethanol. Air-dried pellets were re-suspended in 4.3 mL of Tris-EDTA. After RNAse treatment, nucleic acids were extracted using Phenol chloroform. DNA precipitation was done using Ammonium Acetate 7.5M and absolute ethanol. Each sample was re-suspended in very low Tris-EDTA buffer.

### Illumina sequencing and variant calling procedure

Genomic DNA was sequenced via the Illumina HiSeq 2000 (Genewiz, Leipzig). Short sequence reads were trimmed using Trimmomatic v0.32 (Bolger et al., 2014) and aligned to the masked reference genome (Goodwin et al., 2011), using BWA v0.7.15 (Li et al., 2009). Alignments were converted to BAM using SAMtools v0.1.4 (Li et al., 2009). The percentage of IPO323 chromosomes covered by reads for each sample was calculated via BEDtools genomecov (Quinlan and Hall, 2010). In order to identify SNPs and short indels segregating among the different lineages used in this study, we extracted variants from each sample using the GATK v-3.3-1-0 (McKenna et al., 2010) HaplotypeCaller with the following parameters: -T genotypeGVCFs GVCF -- variant_index_type LINEAR max_alternated_alleles 4-ploidy 1. Identified SNPs and short indels were filtered using VariationFiltration-filterExpression QD < 2.0 || FS > 60.0 || MQ < 40.0. The impact of each variant on annotated genes was estimated using SnpEff v 4.3 (Cingoolani et al., 2012), and variant effects on the secretome were obtained using SnpSift v4.3 (Cingoolani et al., 2012) with a bed file from Morais Do Amaral et al., 2012.

### Statistics

To estimate the significance of symptom differences among the fungal lineages we analyzed symptoms independently for each wheat cultivar. We used the following mixed linear model with the R package *nlme:*

D ~ Bi + Sj + Bi*Sj *+ RegREP(Vir(L)),*

where D is the disease symptoms, Bi is the fixed effects of the genetic background (i = MGGP01 or MGGP44), Sj is the fixed effect of the evolutionary status of fungal samples (j= ancestral, evolved at 17°C, evolved at 23°C or evolved under fluctuations), Bi*Sj the interaction term. For each genetic background, the three lineages per selection regime were included (see **Figure 1**). This level of replication was considered random (*RegREP*), and both virulence assay (Vir) and leaf (L) were nested ramdom effects within the regime replicate effect. For each genetic background, we calculated the contrasts between the evolutionary status *Sj* effect using mutltcomp R package and FDR threshold at 5% (Benjamini-Hochberg, 1995).

## Results

### Quantitative variation of disease symptoms among wheat cultivars

Knowledge on the genetic basis of resistance level among wheat cultivars is incomplete (Brown et al., 2015). In the present panel comprising six bread wheat cultivars, only 4 resistance genes have been identified so far: Stb4 and Stb11 in Apache (Ghaffari et al. 2011), Stb6 in Caphorn (Zhong et al. 2017), and Stb9 in Courtot (Chartrain et al., 2009) (**Figure 2B**). Although disease symptoms were widely scattered (as shown in the boxplots for ancestors MGGP01 and MGGP44 on the cultivar Premio), this study still identified substantial differences in symptom quantity between the ancestors and the evolved lineages.

### Significant decrease of virulence of evolved lineages

In the in vitro evolutionary regimes, the significant effects we observed were solely linked to diminished symptoms. The reduction in symptoms was greatly influenced by both the wheat cultivar and the fungal genetic background (MGGP01 or MGGP44). Figure 3 depicts the distribution of symptoms for each cultivar individually. When comparing the two stable evolution regimes, more significant loss of symptoms was observed following evolution at 17°C compared to that after evolution at 23°C (7 out 10 comparisons) (**Table 1**). While not statistically significant, two out of the three remaining comparisons showed a comparable trend (P > 0.1) between the effective strategies in the tested regimes. The same trend was observed when comparing symptoms between the fluctuation and 23°C selection regimes. In 8 out of 10 comparisons, a greater reduction in symptoms was observed after evolution under fluctuating conditions compared to evolution at 23°C (**Table 1**). The findings indicate that exposure to both 17°C and fluctuating temperatures (ranging between 17°C and 23°C) had a more pronounced impact on disease levels compared to a constant temperature of 23°C. Furthermore, although the fluctuating selection regime rarely resulted in significantly fewer symptoms at 17°C (only three cases, all associated with the genetic background MGGP01), we did not observe the reverse pattern.

**Figure 3 -.**
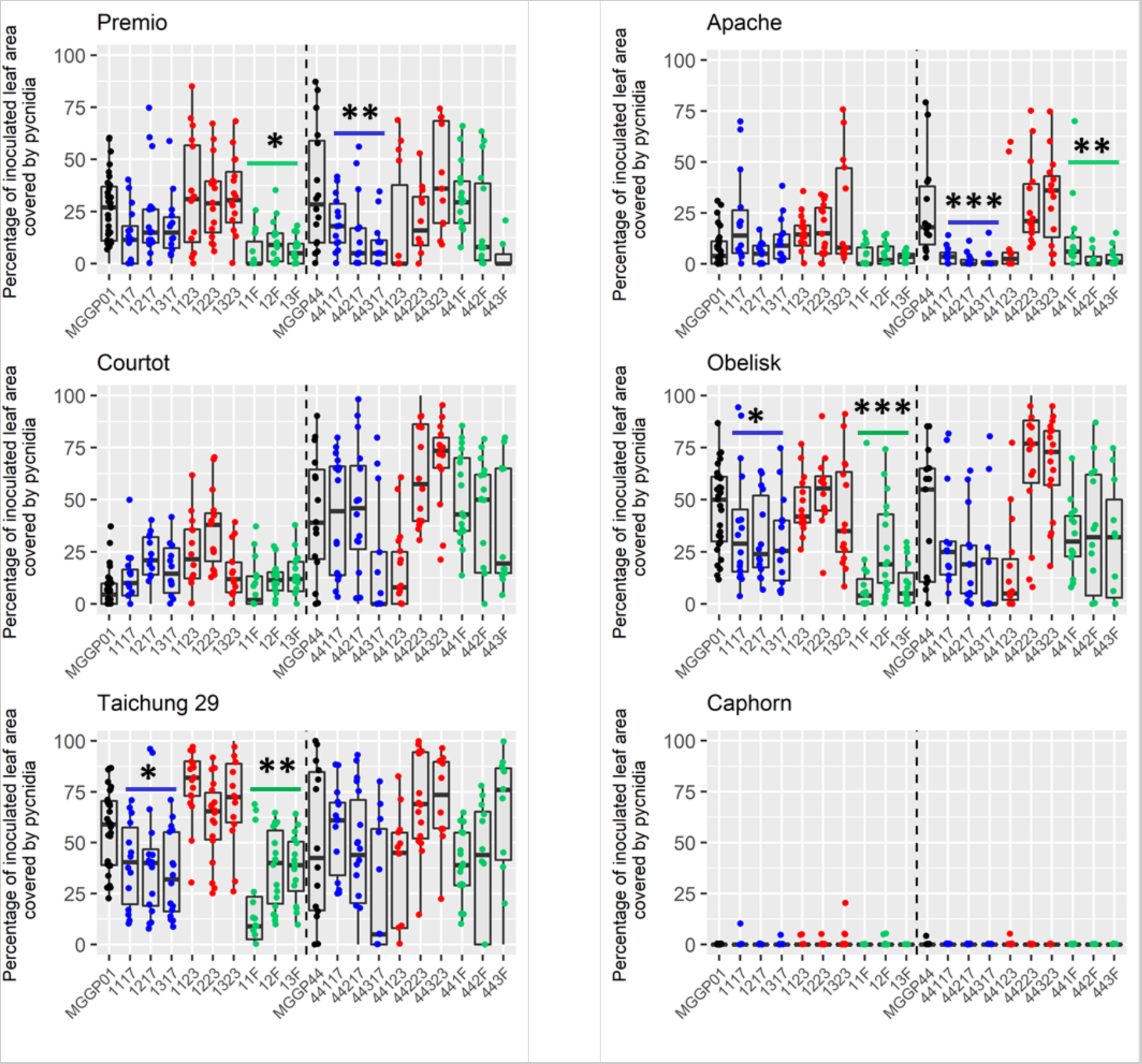
Symptoms for each wheat cultivar-fungal genotype interaction. Boxplots of the distribution of the percentage of leaf area covered with pycnidia (PLACP) is shown for each wheat-fungus interaction. Selection regimes are indicated by color: ancestors (black); Stable 17°C (blue); Stable 23°C (red); fluctuating (green). Significance level of symptom changes in evolved lineages compared to their ancestor are indicated: * (p<0.05); ** (p<0.1), *** (p<0.01). The vertical dashed line separates lineages of MGGP01 genetic background (*left*) and MGGP44 genetic background (*right*).

**Table 1 -.** Comparisons of symptom development between the lineages exposed to different selection regimes. Three selection regimes were used during the experimental evolution: Stable 17°C, Stable 23°C and fluctuating temperature. All comparisons for each genetic background are presented and were tested independently for each wheat cultivars (Premio, Apache, Courtot, Obelisk, Taichung29). FDR adjusted p-values below the 5% threshold are indicated in bold.

|  | 23°C vs 17°C |  | Fluctuations vs 23°C |  | Fluctuations vs 17°C |  |
| --- | --- | --- | --- | --- | --- | --- |
|  | *MGGP01* | *MGGP44* | *MGGP01* | *MGGP44* | *MGGP01* | *MGGP44* |
| Premio | **3.9 x 10<sup>-4</sup>** | **0.027** | **4.7 x 10<sup>-10</sup>** | 0.49 | **0.021** | 0.11 |
| Apache | 0.10 | **2.2 x 10<sup>-10</sup>** | **3.2 x 10<sup>-5</sup>** | **8.1 x 10<sup>-7</sup>** | **0.018** | 0.27 |
| Courtot | 0.39 | 0.090 | **0.038** | 0.65 | 0.44 | 0.39 |
| Obelisk | **6.0 x 10<sup>-3</sup>** | **1.2 x 10<sup>-6</sup>** | **3.3 x 10<sup>-10</sup>** | **8.8 x 10<sup>-4</sup>** | **7.1 x 10<sup>-4</sup>** | 0.17 |
| Taichung | **2.2 x 10<sup>-9</sup>** | **6.0 x 10<sup>-3</sup>** | **3.0 x 10<sup>-13</sup>** | **0.021** | 0.37 | 0.64 |

### No loss of accessory chromosomes

We conducted genome-wide sequencing to verify any changes in the number of accessory chromosomes following the in vitro experimental evolution. Although we observed high level of polymorphism between the two genetic backgrounds, our findings indicate that there is no loss of accessory chromosomes among the sequenced lineages. (**Figure 4**).

**Figure 4 -.**
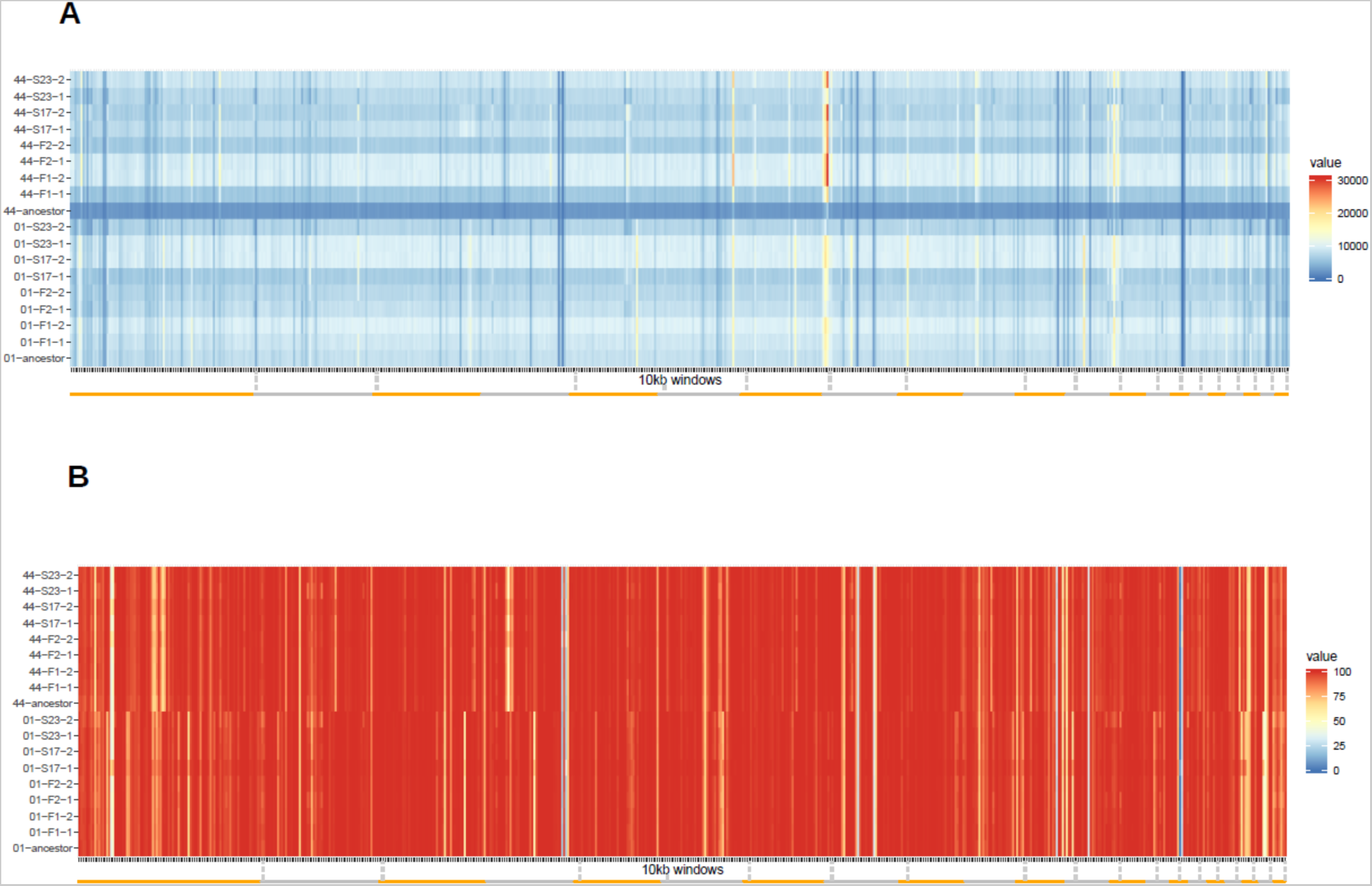
Whole genome sequencing confirms the presence of 21 chromosomes for all lineages. Coverage heatmaps for all samples showing most of coverage differences are between the two ancestral strains MGGP01 and MGGP44 and the 8 evolved lineages grown at 17°C and 23°C. **A.** Average coverage along the genome (10kb windows) **B.** Percentage of very low coverage (<2X) along the genome (10kb windows). Orange and grey bars under each graph are delimiting the 21 chromosomes.

### Pattern of mutation accumulation between the evolved lineages

We detected an average nucleotide diversity of 0.06% across the genome within the evolved lineages (**Table 2**). We still have to be cautious with the interpretation of these results. It is quite probable that the SNP calling method we employed may be overestimating the actual count of mutations occurring during our lab-based experimental evolution. We hypothesize that this discrepancy might be attributed to copy number variants, contributing to an erroneous estimation. We anticipate that long-read sequencing will yield precise variant calls for the evolved genomes. Likewise, further validation of variant calling using Sanger sequencing for a subset of the mutations will be helpful to measure accuracy rate. However, the distribution of SNPs throughout the genome exhibits a distinctive and recurrent pattern among genotypes and regimes, for the most part (**Figure 5A**). For small insertion-deletions, the distribution of variable sites appeared to be random (**Figure 5B**). Employing more tools to concurrently examine small-scale and large-scale mutations will offer deeper insights into the nature and distribution of accumulated mutations throughout the experimental evolution

**Figure 5 -.**
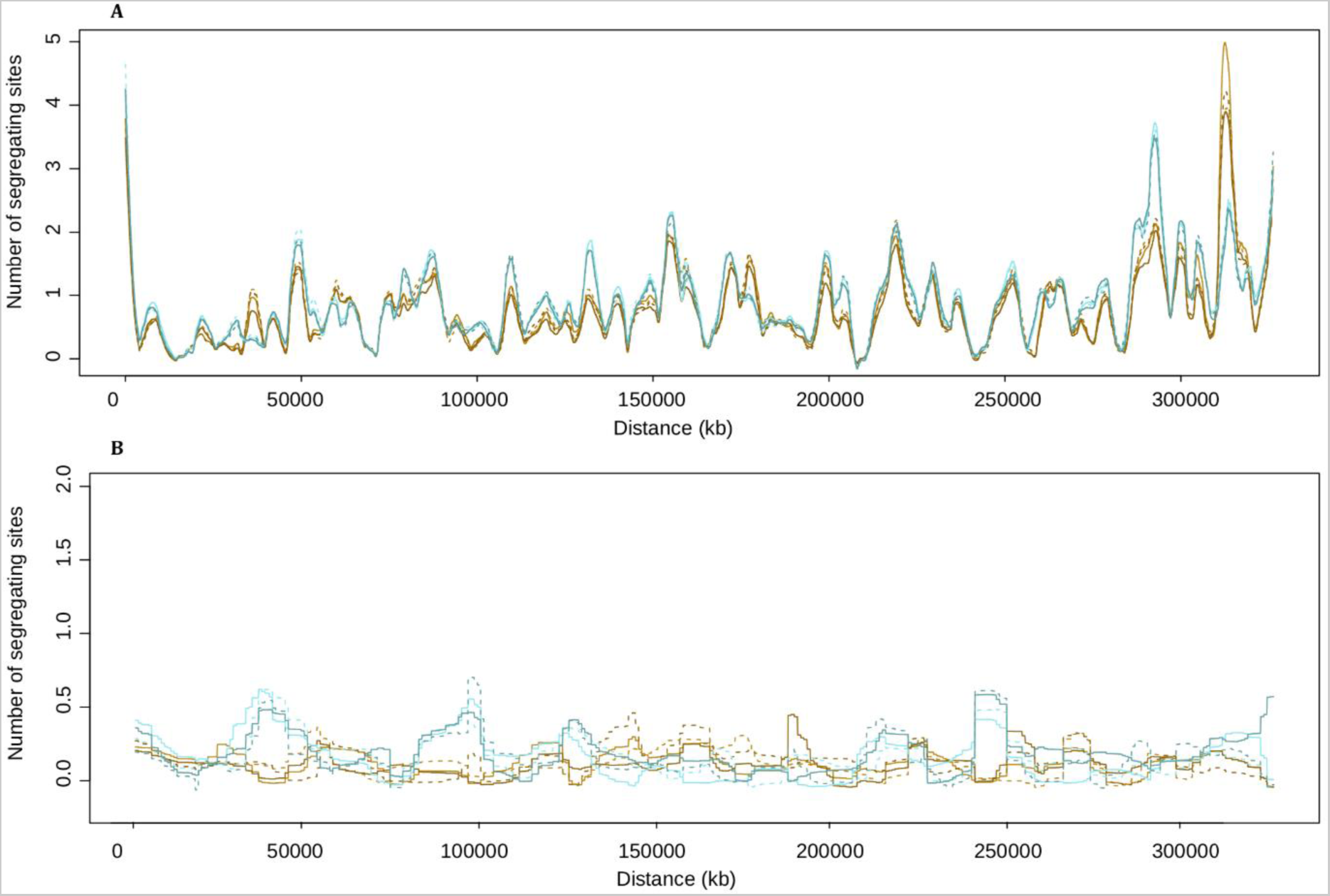
Number of variable SNPs and short insertion deletions in 10kb sliding windows, with an overlap of 100bp. In total 8 groups of samples are represented: in brown, dashed brown, dark brown and dashed dark brown are for Stable 17°C + ancestor, Stable23°C + ancestor, Fluctuating-1+ancestor and Fluctuating-2 + ancestor, for background MGGP01; the same palette but blue was used for the lineages from the genetic background MGGP44. Note that the presence of the ancestor in each group allows us to identify SNPs that could be monomorphic among all evolved lineages but different from their ancestors. **A**: SNPs; **B**: short indels.

**Table 2 -.** Total number of small scale mutations identified per evolved lineage.

| MGGP01 |  |  |  | MGGP44 |  |  |  |
| --- | --- | --- | --- | --- | --- | --- | --- |
| 17°C | 23°C | F-1 | F-2 | 17°C | 23°C | F-1 | F-2 |
| 26011 | 25923 | 23617 | 25420 | 29820 | 29974 | 28983 | 29319 |

### Effect of mutation accumulation on the secretome

By using the predicted secretome annotation established by Morais Do Amaral et al. (2012), we pinpointed mutations within genes encoding proteins predicted to play a role in counteracting the host immune response. Our findings show a limited number of mutations accumulated in secretome genes, dispersed throughout the genome. **Figure 6** illustrates the total count of these mutations categorized by their functional impact on nucleotide substitution, either located within exons (synonymous or non-synonymous), introns, 5 prime- and 3 prime-untranslated regions (UTRs) (no stop codon mutation were detected) (**Figure 6A** and **6B**). We detected a large number of non-synonymous mutations for both genetic backgrounds (**Figure 6 A** and **B**) and also a large number of mutations in introns for the background MGGP44. Effector genes are anticipated to accumulate variants due to positive or diversifying selection driven by the selective pressure imposed by the host immune system. Contrary to expectations, the discovery of a higher count of non-synonymous mutations compared to synonymous ones after in vitro evolution is rather surprising. Additionally, we present the mutational effects of small insertions and deletions identified within this particular set of coding regions (**Figure 6C** and **6D**). We detected a few disruptive effects of indels. For both backgrounds, we found an excess of intronic and frameshift mutations. To confirm the nature and precise location of these variants, long-read sequencing is necessary. This will provide a more thorough examination to determine whether these random mutations contribute to adaptive evolution against the host immune system.

**Figure 6 -.**
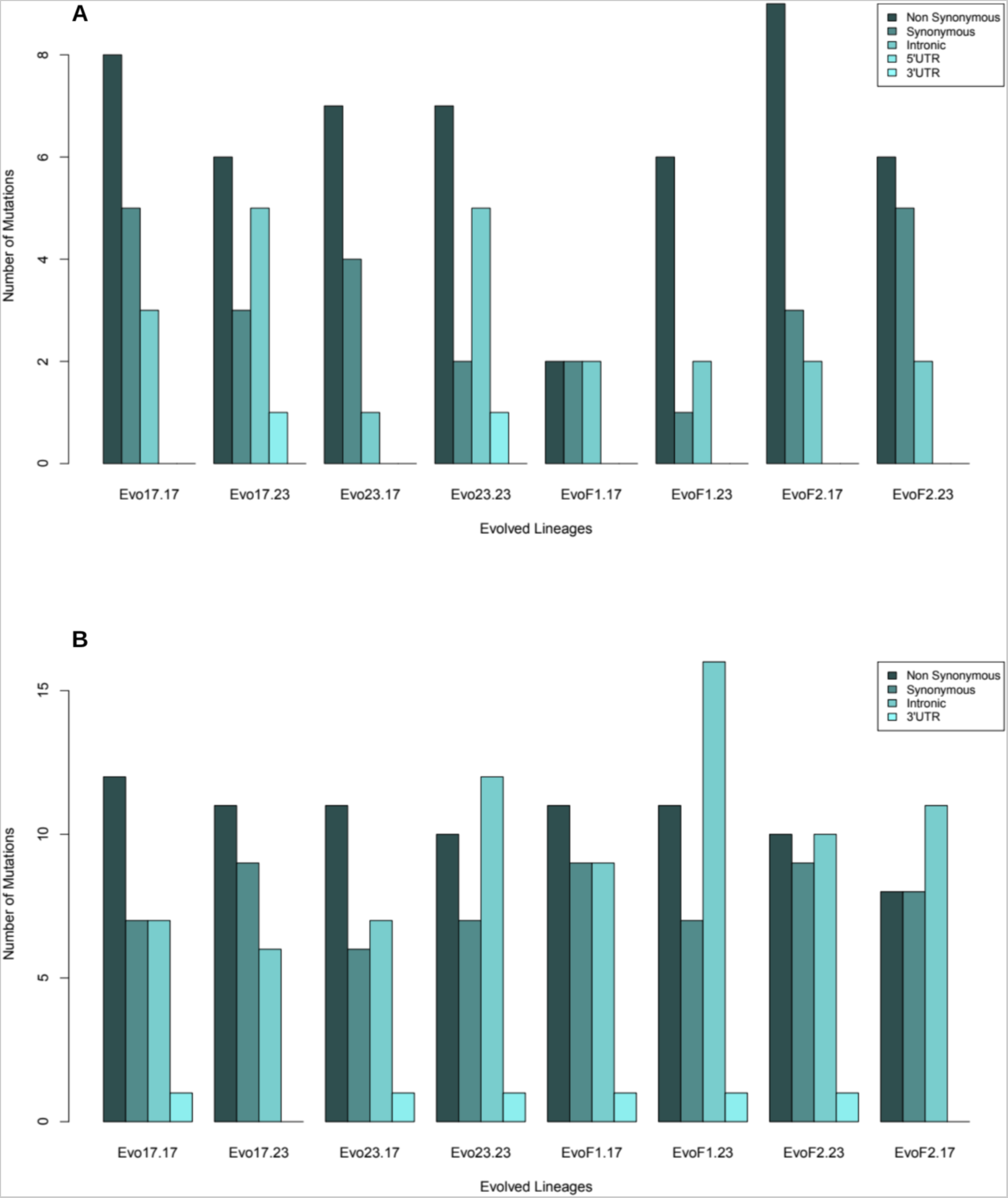
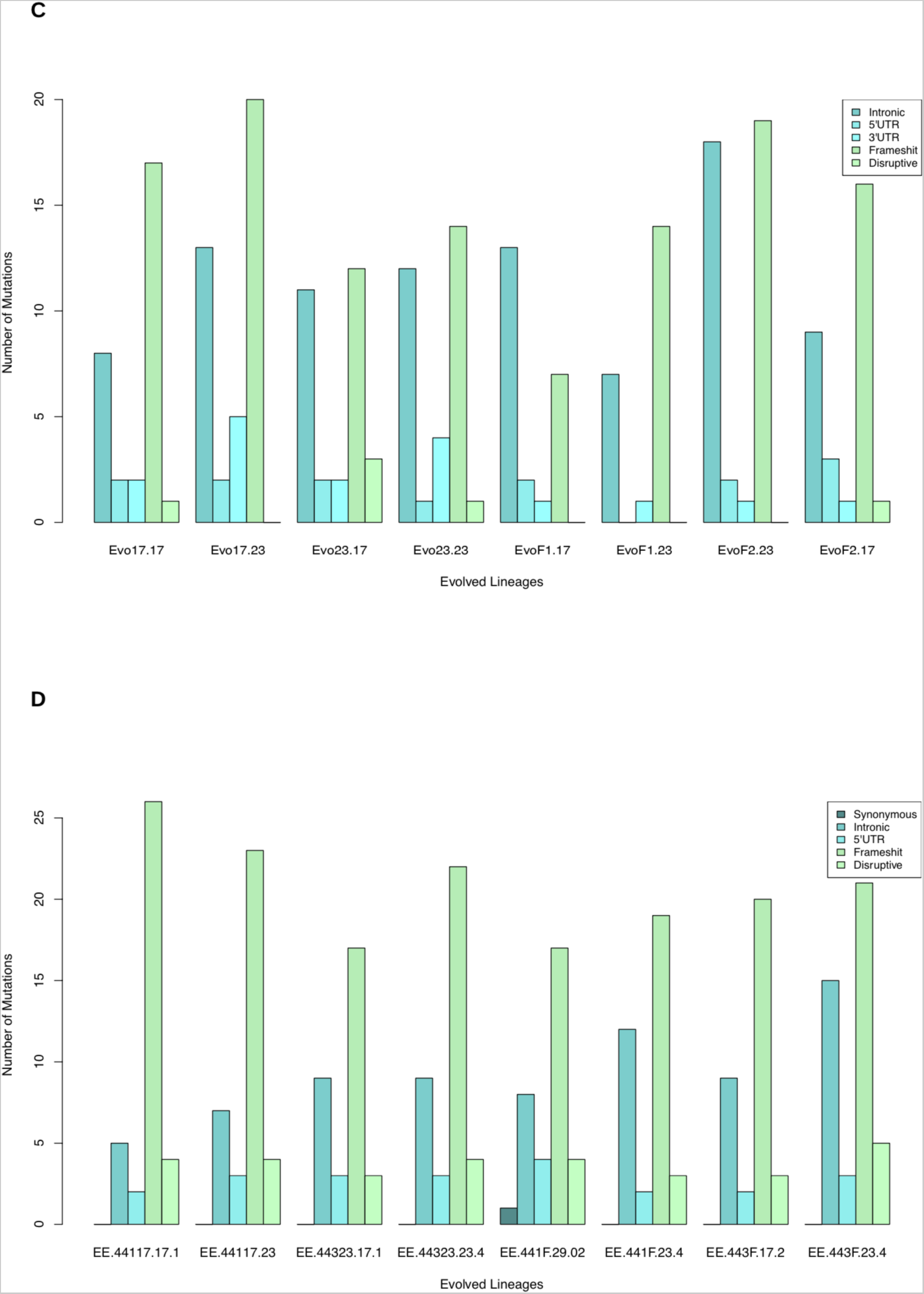
Number of mutations in genes annotated in Z. tritici secretome for the genetic background MGGP01 and MGGP44. Categories of variant annotation were identified using snpSift (Cingolani et al., 2012) **A**: SNPs for MGGP01 **B**: SNPs for MGGP44; **C**: indels for MGGP01; **D**: indels for MGGP44. The location and functional consequences of the mutations are color-coded (see Figure legends).

## Discussion

This work represents the first attempt towards understanding the influence of the selection under variable environment on the virulence level of the fungus *Z. tritici,* using experimental evolution. Using a panel of six wheat cultivars which are gradually different for their resistance level against *Z. tritici*, our results support a loss of virulence although the phenotypic response between evolved lineages is often heterogeneous. Our results also suggest that a complex genetic basis is underlying the quantitative variation of pycnidiae produced by the pathogen, in agreement with former studies, also suggesting that many unknown molecular factors, most likely of small or moderate phenotypic effect, contribute to the phenotype (Stewart et al., 2017). Due to the lack of power in our study we did not investigate the link between genotype and phenotype through association tests. However we were still interested to seek new mutations in the evolved lineages that could be of strong relevance regarding to their function. To reach this goal, we annotated all mutations in the secretome, and found a few sites and small indels that were variable between the evolved lineages and their ancestor. The nature of these mutations (such as amino-acid substitutions or mutations disrupting the reading frame of coding regions) underscores the considerable evolutionary potential of this fungal species. Whether these mutations are genuine mutations associated causing a decrease of virulence remains to be elucidated. These new alleles were either unique to a lineage, or shared between lineages. Trans-lineage polymorphism is not an expected result if we consider that mutations accumulated following a random walk during the process of evolution in our experiment.

It is known that many rearrangements occur during meiosis (Croll et al., 2015; Fouché et al., 2018). Our understanding of structural variations (SV) occurring during mitosis is limited, we suggest structural rearrangements driven by TE transpositional activity can still take place during mitoses. Following long read sequencing we could reveal structural rearrangements (SV). These SV are well known in fungal genomes and *Z. tritici* genome to play a role in adaptation. Here the substantial difference in coverage across the genome between the two strains used as ancestors for our experimental evolution further exemplifies the widespread presence of structural variations (SV) within *Z. tritici* populations.

This study specifically targeted genes within the secretome. Nevertheless, numerous other variable sites emerged when comparing the alleles in evolved lineages to those of the ancestors. Some of these mutations might also contribute to the observed decrease in disease among the evolved lineages; we cannot rule out their potential impactFirstly, it’s probable that not all effector proteins have been annotated yet, and this includes genes that might exist in our two ancestors but are absent from the reference genome (Plissonneau et al., 2018). Secondly, mutations in genes outside of the secretome could also impact the level of virulence. Whether these mutations affect regulatory or coding regions, targeting effector genes or others, it hints at potential antagonistic pleiotropy. Some genes might contribute both to in vitro growth and the pathogen’s development *in planta.* Significantly, a previous QTL study on Z. tritici pinpointed several candidate genes exhibiting pleiotropic effects, influencing both melanization and morphology (Lendenmann et al., 2016).

These findings underscore the significant role of the abiotic environment in shaping the evolution of the Z. tritici pathogen. Hence, it becomes crucial to account for these factors in devising management strategies aimed at controlling pathogen populations

